# The chick caudo-lateral epiblast acts as a permissive niche for generating neuromesodermal progenitor behaviours

**DOI:** 10.1101/243980

**Authors:** Peter Baillie-Johnson, Octavian Voiculescu, Penny Hayward, Benjamin Steventon

## Abstract

Neuromesodermal progenitors (NMps) are a population of bipotent progenitors that maintain competence to generate both spinal cord and paraxial mesoderm throughout the elongation of the posterior body axis. Recent studies have generated populations of NMp-like cells in culture and have been shown to differentiate to both neural and mesodermal cell fates when transplanted into either mouse or chick embryos. Here, we aim to compare the potential of mouse embryonic stem (ES) cell-derived progenitor populations to generate NMp behavior against both undifferentiated and differentiated populations. We define NMp behaviour as the ability of cells to i) contribute to a significant proportion of the anterior-posterior body axis, ii) enter into both posterior neural and somitic compartments and, iii) retain a proportion of the progenitor population within the posterior growth zone. We compare previously identified ES cell-derived NMp-like populations to undifferentiated mouse ES cells and find that they all display similar potentials to generate NMp behaviour *in vivo*. To assess whether this competence is lost upon further differentiation, we generated anterior and posterior embryonic cell types through the generation of 3D gastruloids and show that NMp competence is lost within the anterior (Brachyury negative) portion of the gastruloid. Taken together, this demonstrates that the chick caudo-lateral epiblast is a highly permissive environment for testing NMp competence and is therefore not suitable as a positive test of neuromesodermal progenitor identity *(ie*. specification). However, it does act as an appropriate system to test for the loss of NMp potential, and therefore offers insight as a functional test for the regulation of NMp competence *in vivo*.

## Introduction

Amniotes form the posterior tissues of their bodies progressively, from populations of progenitors cells situated within the posterior growth zone. Neuromesodermal progenitors (NMps) are one such population of posterior progenitors that continuously allocate cells to both the spinal cord and presomitic mesoderm (PSM) (Selleck and Stern, 1991; Brown and Storey, 2000; Tzouanacou *et al*., 2009; Henrique *et al*., 2015). This ability to maintain multi-germ layer competence past primary gastrulation stages presents a unique opportunity to understand the molecular mechanisms that maintain cellular competence during development (Steventon and Arias, 2016). However, the degree to which this is an autonomous property of the cells, or else a non-cell autonomous property given by their signaling environment, is unknown.

The functional properties of the embryonic NMp population have been demonstrated through transplantation experiments (Cambray and Wilson, 2002, 2007) and retrospective clonal analyses (Tzouanacou *et al*., 2009). The former have demonstrated their capacity not only to contribute to neural and mesodermal tissues across many levels of the body axis, but also their ability to self-renew through serial heterochronic transplantation (Cambray and Wilson, 2002). Embryonic NMps can therefore maintain their competence to form both neural and mesodermal tissues beyond the normal duration of somitogenesis, when introduced into a permissive niche such as that found in the earlier embryos. The common origin for these tissues was demonstrated conclusively through clonal labelling experiments that indicated the continued presence of a bipotent population throughout axial elongation in the mouse (Tzouanacou *et al*., 2009). The patterns of clonal labelling predicted phases of expansion and depletion that have been confirmed subsequently by quantification of the number of Sox2^+^, T/Brachyury^+^ cells during axial elongation (Wymeersch *et al*., 2016).

The signals and transcription factors expressed in this embryonic domain are key features of protocols used to generate and maintain Sox2 and T/Brachyury co-expressing cells *in vitro* (Gouti *et al*., 2014; Tsakiridis *et al*., 2014; Turner *et al*., 2014). Treatment with the canonical Wnt signaling agonist CHIR99021 (Chiron) and exogenous FGFs are common features of these protocols. Tsakiridis *et al*. describe how, in cultures of mouse epiblast stem cells, the pulse of Wnt signalling acts to generate a mixture of committed mesendodermal and neuromesodermal progenitors from an initially primitive-streak biased subpopulation of the original culture (Tsakiridis *et al*., 2014). The pulse of Chiron upregulates expression of Brachyury, as well as Hox genes from paralogous groups 5-9, which are active late in gastrulation. Similar results are obtained by Gouti *et al*., who describe the formation of Brachyury and Sox2 co-expressing cells in cultures of mouse embryonic stem cells that have been grown in FGF2 and treated with a pulse of Wnt signalling (Gouti *et al*., 2014, 2017). We have shown that a pulse of Chiron, acting in concert with endogenous FGF signalling, can direct adherent cultures of mouse embryonic stem cells towards a neuro-mesodermal progenitor-like state (Turner *et al*., 2014, 2017). However, expression of Sox2 and Brachyury is often accompanied by the expression of multiple other tissue specific markers within these progenitor populations, suggesting that they resemble a mixed population of multipotent progenitors rather than a pure population of NMps (Edri *et al*., 2018).

In the absence of a definitive gene expression signature, transplantation to the mouse or chicken embryo has been used as a functional test of the capacity of *in* vitro-differentiated cells to produce NMp-like contributions, similar to those described for the embryonic population (Cambray and Wilson, 2002, 2007; McGrew *et al*., 2008). Transplantation into the mouse embryo is technically demanding due to its small size, its initially cup-shaped geometry and the internal nature of its development. Live imaging the incorporation of grafted cells adds an additional layer of complexity, so previous transplantations of *in vitro*-derived cells have been limited to endpoint analyses such as that reported by Tsakiridis *et al*. (Tsakiridis *et al*., 2014). The chicken embryo offers a more accessible host system for testing the *in vivo* potential of candidate NMp populations due to its large size and its external development. Gouti *et al*., showed that the grafted cells incorporate into both the neural tube and the somitic compartments (Gouti *et al*., 2014). While these experiments demonstrate the competence of *in* vitro-derived Sox2^+^, T/Brachyury^+^ populations to generate both neural and mesodermal derivatives *in vitro*, it is not clear whether this is a specific property of NMp-specified populations, or else a consequence of a multipotent population of cells responding to transplantation into a permissive niche. To address this, we set out to compare the ability of these cells to generate NMp behaviour to grafts of either uncommitted or more differentiated ES cells when transplanted into the chick caudo-lateral epiblast (CLE).

## Results and Discussion

### The Region Lateral to the Node Contributes Extensively to Both Neural and Mesodermal Tissue

To identify a region of the embryo in which to graft candidate populations of neuro-mesodermal progenitors, a fate map was constructed by labelling small areas of the ectoderm with Dil (see Materials and Methods). The position of the labelled cells was recorded in relation to the caudal limit of the node after labelling and the size of the label was measured in the rostro-caudal (hereafter “axial”) direction. After 15 hours’ incubation, the distance between the most rostral and most caudal labelled cells was measured and is shown in Figure 1C (and Supplementary Figure 2A). The contribution of the labelled cells to the different tissue compartments was determined (Supplementary Figure 2B) and is summarised in Figure 1D. On mapping the labels in this way, it is clear that labelled cells positioned in a small region lateral and slightly caudal to the node (shown as a dashed box in Figures 1-3 and Supplementary Figures 2-9) can contribute to extensively to both neural and mesodermal tissues (a “mixed” contribution), as is consistent with previously published fate maps of this region (Selleck and Stern, 1991; Catala *et al*., 1996; Brown and Storey, 2000). In order to discriminate between contributions of different lengths, a series of thresholds were used: a lower one at 500μm, which is less than the length of three somites and an upper threshold of 1750μm (see Supplementary Figure 1). Where the distance between the most rostral and most caudal labelled cells was less than the lower threshold, the contribution was defined as “short”; between the thresholds was defined as “medium” and above the upper threshold was defined as “long”.

**Figure 1:**
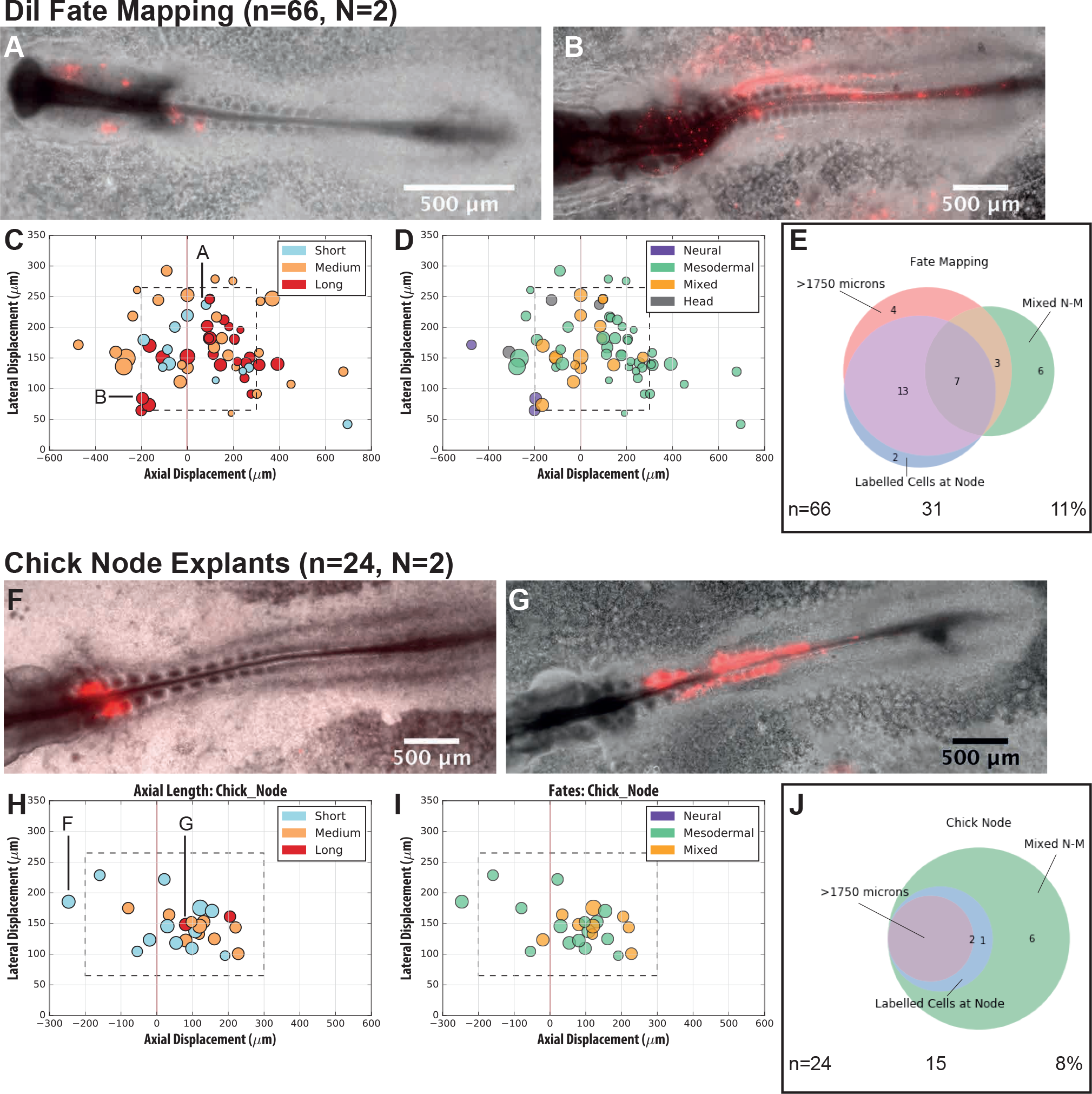
Embryonic Cells Caudal and Lateral to the Node Show NMp-like Behaviour and are Sampled in Chicken Node Explants. The location of the embryonic NMp population lateral to the node was determined through Dil tissue labelling (A-E) and confirmed by heterotopic transplantation of explanted node tissue (F-J). The starting position of each label was measured as the absolute axial and lateral displacement from the caudal limit of the node (the x and y axes, respectively, in (C), (D), (H) and (I); node at (0,0)). The length of the rostro-caudal spread of labelled cells was measured on each side of the midline and their contribution to the different tissue compartments was scored by inspection. (A/F) and (B/G) show the shortest and longest labelled cell contributions, respectively. (C/H) shows the starting position of each label, coloured according to the final length of the labelled cell contribution: <500μm, blue; 500–1750μm, orange and >1750μm, red. (D/I) shows the starting position of each label, coloured according to whether the labelled cells contributed to neural tissues (purple), mesodermal tissues (green) or both (orange). Contributions to the anterior tissues of the head are indicated in grey. A rectangular region of interest (dashed line in (C), (D), (H) and (I)) is defined as the largest region that encompasses all of the long, mixed neural and mesodermal contributions. The Venn diagrams in (E) and (J) summarise the overall behaviour of the labels or grafts. Each label/graft was scored into the three sets, corresponding to long contributions, mixed neural and mesodermal contributions and cases where labelled cells resided in the region around the node. The number of single-germ layer restricted, short or medium labels/grafts that did not occupy the region around the node is indicated. n denotes the number of labels/grafts; N the number of biological replicates. Scale bars indicate 500μm throughout.

**Figure 2:**
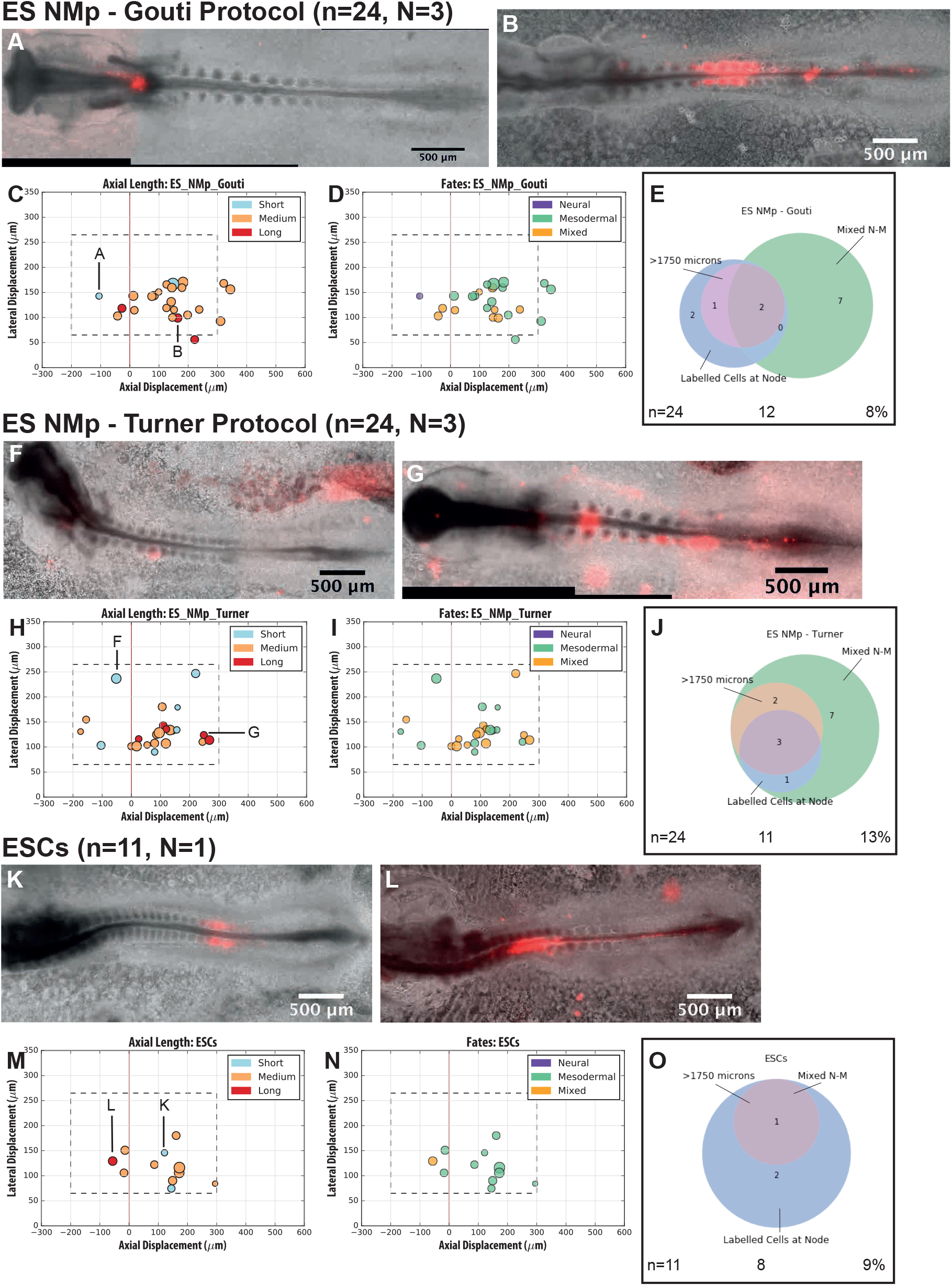
Embryonic Stem Cells (ESCs) and ESC-derived NMps are Competent to Produce NMP-like Contributions on Transplantation. Labelled embryonic stem cells (ESCs, (K-O)) and ESC-derived NMps ((A-E) and (F-J)) were transplanted into the region caudal and lateral to the node. The starting position of each graft was measured as the absolute axial and lateral displacement from the caudal limit of the node (the x and y axes, respectively, in (C), (D), (H), (I), (M) and (N); node at (0,0)). The length of the rostro-caudal spread of labelled cells was measured on each side of the midline and their contribution to the different tissue compartments was scored by inspection. (A/F/K) and (B/G/L) show the shortest and longest labelled cell contributions in each case, respectively. (C/H/M) shows the starting position of each graft, coloured according to the final length of the labelled cell contribution: <500μm, blue; 500–1750μm, orange and >1750μm, red. (D/I/N) shows the starting position of each graft, coloured according to whether the labelled cells contributed to neural tissues (purple), mesodermal tissues (green) or both (orange). The Venn diagrams in (E), (J) and (O) summarise the overall behaviour of the grafts. Each graft was scored into the three sets, corresponding to long contributions, mixed neural and mesodermal contributions and cases where labelled cells resided in the region around the node. The number of single-germ layer restricted, short or medium grafts that did not occupy the region around the node is indicated. n denotes the number of grafts; N the number of biological replicates. Scale bars indicate 500μm throughout.

**Figure 3:**
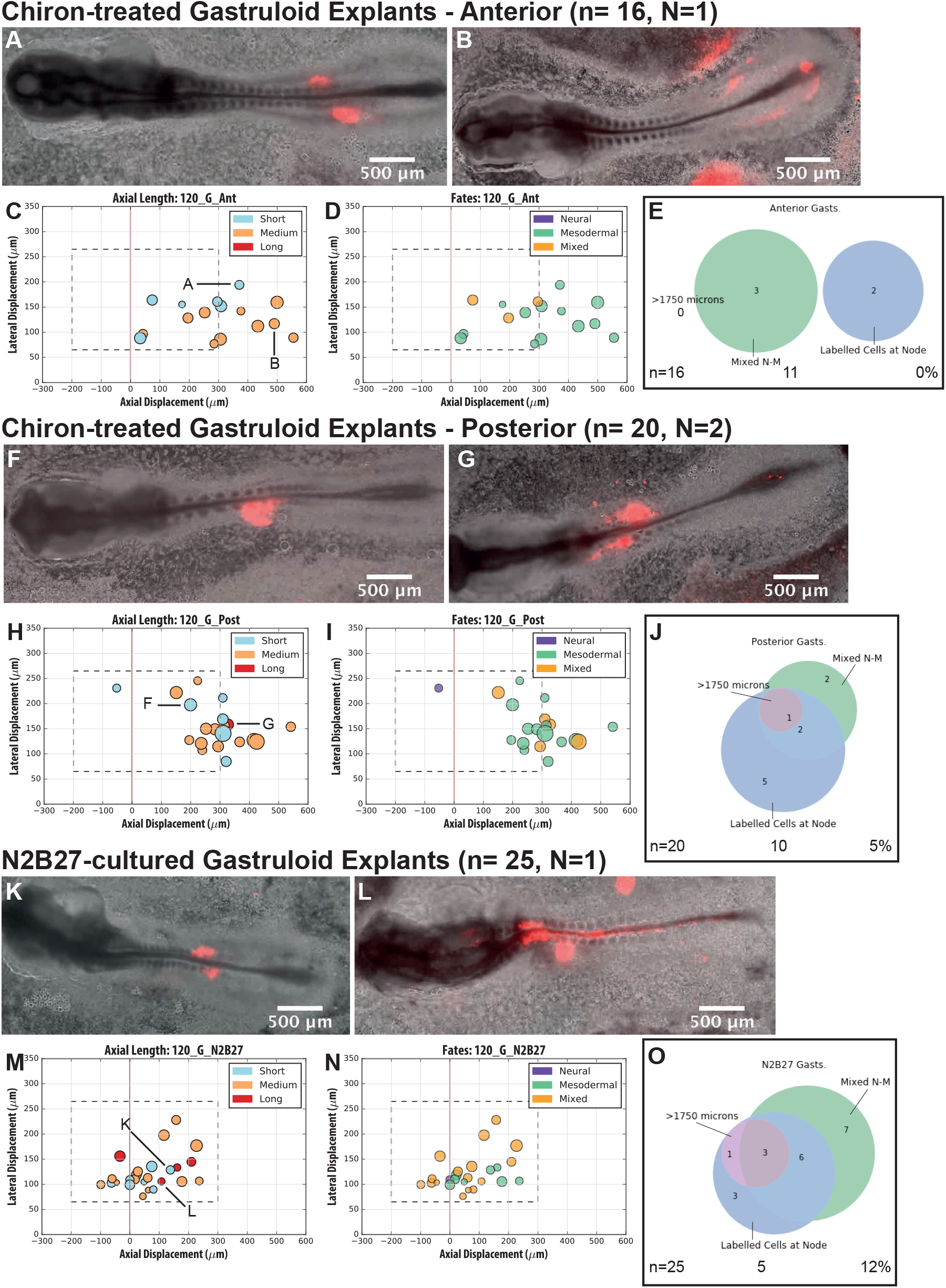
Anterior Explants from Chiron-treated Gastruloids are Not Competent to Produce NMp-like Behaviour, unlike Posterior or Untreated Explants. Labelled tissue explants from Chiron-treated (anterior, (A-E)) or posterior (F-J)) or untreated (K-O) gastruloids were grafted into the region caudal and lateral to the node. The starting position of each graft was measured as the absolute axial and lateral displacement from the caudal limit of the node (the x and y axes, respectively, in (C), (D), (H), (I), (M) and (N); node at (0,0)). The length of the rostro-caudal spread of labelled cells was measured on each side of the midline and their contribution to the different tissue compartments was scored by inspection. (A/F/K) and (B/G/L) show the shortest and longest labelled cell contributions in each case, respectively. (C/H/M) shows the starting position of each graft, coloured according to the final length of the labelled cell contribution: <500μm, blue; 500–1750μm, orange and >1750μm, red. (D/I/N) shows the starting position of each graft, coloured according to whether the labelled cells contributed to neural tissues (purple), mesodermal tissues (green) or both (orange). The Venn diagrams in (E), (J) and (O) summarise the overall behaviour of the grafts. Each graft was scored into the three sets, corresponding to long contributions, mixed neural and mesodermal contributions and cases where labelled cells resided in the region around the node. The number of single-germ layer restricted, short or medium grafts that did not occupy the region around the node is indicated. n denotes the number of grafts; N the number of biological replicates. Scale bars indicate 500μm throughout.

By combining the information on the length of the labelled cells’ contributions and the tissues they occupied, it was possible to define a square region of interest (ROI) lateral to the node, which included all of the long contributions to both neural and mesodermal tissues. It was noted that some of the long, mixed contributions also included cells resident in the region around the node at the end of the incubation period (red points in Supplementary Figure 2). When cells were labelled rostrally to this ROI, they could contribute exclusively to neural tissues, while labels placed more caudally were exclusively mesodermal. The results of the fate mapping therefore define a region in which to graft populations of candidate NMps in order to test their capacity to contribute to neural and mesodermal tissues across many axial levels.

As as a positive control for the generation of NMps in this assay, we labelled explants of the region around and including the chicken node and grafted them into the region defined by the fate mapping experiment. When grafted into the ROI, the explants were found to give mostly mesodermal and mixed contributions that were infrequently long (see Figure 1F-J and Supplementary Figure 3). Time lapse imaging showed that the long contributions arose through labelled cells remaining in the region around the node as it regressed, with some cells leaving to occupy the presomitic mesoderm (PSM). The results of grafting the node explants show that a candidate neuro-mesodermal progenitor population would be expected to mimic this pattern of NMp-like behaviour in producing long mixed contributions at an equal or higher frequency (_≥_8.70%, with the number of grafts within the NMp-fated domain being n_ROI_=23).

### Embryonic Stem (ES) Cells and ES Cell-derived Candidate NMps are Competent to Show NMp-like Behaviour on Transplantation

To further test the suitability of this assay for testing murine ES cell-derived NMps, candidate cells were produced in accordance with published protocols and were transplanted to the region caudal and lateral to the node. Cells cultured according to the protocol used by Gouti *et al*. have previously been described to contribute to both neural and mesodermal tissues (Gouti *et al*., 2014). In this assay, these cells were found to produce mostly mixed and mesodermal contributions, some of which were long (see Figure 2A-E and Supplementary Figure 6). The frequency of long, mixed contributions within the ROI was found to be 10.00% (n_ROI_=20). Grafts were also observed to contribute labelled cells to the PSM and to the region around the node.

Candidate NMps generated with the protocol published by Turner *et al*. have not previously been tested *in* vivo. In this assay, they were found to give mostly mixed and mesodermal contributions, some of which were long (see Figure 2F-J and Supplementary Figure 5). The long, mixed contributions occurred at a frequency of 20.83% (n_ROI_=24) and included labelled cells in the region of the node and in the PSM (see Supplementary Figure 5). In a manner similar to the grafted node explants, the labelled cells were observed in time-lapse experiments to transiently reside in the region around the node during its regression. Collectively, these results demonstrate that murine ES cells cultured according to both protocols are competent to show NMp-like behaviour on transplantation to the region caudal and lateral to the node. These observations validate the use of this assay as these populations have been described previously as NMp-competent.

On grafting self-renewing cultures of ES cells, however, a similar pattern of long, neural and mesodermal contribution was observed at a frequency of 9.09% (n_ROI_=11, Figure 2K-O) after transplantation to the same region. These grafts appeared to integrate efficiently into the embryonic tissues, with every graft appearing to spread in some way during the incubation. One graft was observed to produce a long, mixed contribution that included labelled cells in the region around the node (Figure 2L and Supplementary Figure 4), while the remaining ten grafts contributed to limited extents of the mesodermal tissues. As a culture of pluripotent cells, it is not surprising that the colonies were competent to make continued contributions to both neural and mesodermal tissues when grafted into a region that is permissive for the embryonic progenitor population. This does, however, raise a cautionary note on the use of transplantation as a functional test of candidate NMp populations when undifferentiated cells retain a capacity to give NMp-like contributions. Although this assay allows this behaviour to be demonstrated for a given population, it does not define a population of NMps as a specific cell type, as many populations may be competent to behave in the same way when transplanted to this region.

### Anterior Explants from Chiron-treated Gastruloids are Not Competent to Produce NMp-like Behaviour, unlike Posterior or Untreated Explants

In order to test whether this assay allows cells that are not competent to show NMp-like behaviour to be identified, more differentiated cultures were transplanted to the region caudal and lateral to the node. Three-dimensional gastruloid cultures were selected for this purpose, due to the fact that they develop a clear antero-posterior polarity (Turner *et al*., 2017) with a candidate NMp population thought to reside in the more posterior tissues (Turner *et al*., 2014). The anterior tissues are thought to be mostly mesodermal and locally express genes such as GATA6, thought to indicate a population of precardiac mesoderm (Turner *et al*., 2017). The antero-posterior polarity of Chiron-treated gastruloids can be scored reliably through phase contrast microscopy alone (Baillie-Johnson, 2017), allowing them to be dissected into corresponding tissue explants prior to transplantation.

On grafting the anterior tissue explants to the ROI, 13/16 of the grafts were observed to give short to medium length mesodermal contributions (Figure 3C), including labelled cells in the PSM and in one case, the region around the node (see Supplementary Figure 7). The remaining three grafts produced mixed contributions, though none of these was long. These tissues clearly lack the competence to produce NMp-like behaviour when grafted into this region of the embryo. In contrast, the posterior gastruloid tissues additionally produced a long, mixed neural and mesodermal contribution, indicating that a subpopulation of these cells retains the competence to show NMp-like behaviour in this assay. Many of the grafts were observed to contribute to the PSM and to the region around the node (see Supplementary Figure 8) at a higher frequency than for the anterior explants (40% versus 12.5%; n=20 and 16, respectively). The frequency of NMp-like contributions appears to be around half of that observed for the chick node explants, ES cells and ES cell-derived NMps, suggesting that the more differentiated gastruloid tissues have more limited competence in this assay.

In order to test this idea further, tissue explants from gastruloids that had not been exposed to Chiron were tested in this assay. Gastruloids cultured in N2B27 alone develop entirely under the influence of endogenous signalling, so likely represent a less differentiated population compared to those treated with Chiron, which promotes symmetry-breaking and axial elongation. On grafting these tissues, the labelled cells produced mostly mixed contributions, which were long at a frequency of 12% (n_ROI_=25; see Figure 3M-O). Time lapse imaging showed that labelled cells often left the main part of the graft to reside in the region around the node as it regressed (52% of 25 cases), though labelled cells were never found in the PSM (see Supplementary Figure 9). Collectively, the frequency of long mixed contributions indicates that these explants behave like the 2D culture-derived candidate neuro-mesodermal progenitor populations. Given the higher frequency of mixed contributions, it appears that this culture produces cells with a greater propensity for neural differentiation than following the Chiron treatment. It therefore seems that the N2B27-cultured gastruloid explants are more frequently competent to show NMp-like behaviour on transplantation to the region caudal and lateral to the node.

In order to test whether the lengths of the axial contributions from the candidate neuro-mesodermal progenitor populations differ from those observed for pluripotent cells, the length measurements were plotted as box and whisker diagrams (Supplementary Figure 10). When presented in this way, it is clear that all of the populations tested, with the exception of the anterior Chiron-treated gastruloids, are capable of producing long, mixed contributions after transplantation (Supplementary Figure 10C). Taken together, these results show that the chick CLE is a permissive environment for competent cells to contribute to neural and mesodermal compartments of the elongating body axis. Under such conditions, derivatives of the grafted populations will be exposed to the appropriate signals for the specification to either cell fate. Therefore, this acts as an appropriate assay to assess the competence of cells to generate neural and mesodermal derivatives *in vivo*. However, we suggest caution in the use of this assay as a positive proof of NMp identity, as it is likely that any multipotent population would generate derivatives that enter into either or both embryonic compartments and differentiate accordingly. This may arise through responses to the signalling environment at the graft site and morphogenetic movements such as tissue flows that also distribute cells along the rostro-caudal axis, giving the impression of a contribution to axial elongation. Nevertheless, the results from this assay show that more differentiated cells (such as the anterior gastruloid explants) cannot be made to behave like NMp-competent populations on grafting to the caudal lateral epiblast.

## Materials and Methods

### Chicken Embryo Culture

Fertilised chicken eggs were stored in a humidified 10°C incubator for up to one week until required. Eggs were transferred to a humidified, rocking 37°C incubator for 24 hours prior to the preparation of embryo cultures.

Embryo cultures were prepared according to a modified version of New Culture (New, 1955) using glass rings with rectangular cross sections to support the explanted embryonic membranes within 35mm diameter bacterial Petri dishes (BD Falcon 351008), on a layer of thin albumen. Pannett-Compton saline was prepared to the formulation as described by Voiculescu *et al*. (Voiculescu, Papanayotou and Stern, 2008). Embryo cultures were incubated in a humidified 37°C incubator prior to use and were staged according to the Hamburger and Hamilton (HH) Series (Hamburger and Hamilton, 1951; reprinted 1993). The embryos typically ranged from HH4–9 after the 24 hour incubation described above. Embryos were freshly fixed in 4% Paraformaldehyde (v/v, in PBS) at the end of the incubation time.

### Fate Mapping

Small quantities of CellTracker™ Red (C34552, Thermo Fisher Scientific; prepared in 20% sucrose (v/v)) were micropipetted onto the ectoderm around the node with a mechanically drawn capillary needle and a mouth pipette (Sigma-Aldrich, A5177), within a small droplet of Pannett-Compton saline. Two sites were labelled per embryo, either side of the midline; the droplet of saline was aspirated immediately after the second label had been placed to remove excess dye.

### Mouse Embryonic Stem Cell Culture

E14Tg2a (Hooper *et al*., 1987), T/Bra::GFP (Fehling *et al*., 2003) or Nodal^HBE-YFP^ (Papanayotou *et al*., 2014) embryonic stem cells were routinely cultured on 0.1% gelatin (Sigma-Aldrich G1890) pre-coated plastic and were maintained in ES+LIF medium (500mL Glasgow’s Minimal Essential Medium (GMEM; Gibco 11710-035), 5mL sodium pyruvate (Invitrogen 11360-039), 5mL non-essential amino acids (Gibco/Invitrogen 11140-035), 5mL GlutaMAX (Gibco 35050-038), 1mL β-mercaptoethanol (Gibco/Invitrogen 31350-010), 50mL foetal bovine serum (FBS, Biosera FB-1090/500), 550μL Leukaemia Inhibitory Factor (Recombinant LIF, produced in-house by the Wellcome Trust – Medical Research Council Cambridge Stem Cell Institute)). Cell cultures were maintained in humidified incubators at 37°C with 5% CO_2_. Two-thirds of the culture medium was replaced daily with fresh, warm ES+LIF and the cultures were passaged into new flasks every two to three days as required by enzymatic dissociation with 0.05% Trypsin-EDTA (Gibco 25300-054). Samples of embryonic stem cells used for grafting were cultured in ES+LIF and were collected two days after plating.

### Candidate NMp Differentiation

Culture media used for differentiation or for gastruloid culture were prepared in a base medium of N2B27 (NDiff 227^®^; Takara Bio, Y40002), supplemented with CHIR99021 (Chiron; Tocris Biosciences, 4423) and/or Fibroblast Growth Factor 2 (FGF2; R&D Systems, 3139-FB) as appropriate. Adherent cultures of candidate NMps were produced using the protocols described by Gouti *et al*. and Turner *et al*. (Gouti *et al*., 2014; Turner *et al*., 2014, respectively) and were collected 72 hours after plating. Gastruloid cultures were prepared as described previously (Turner *et al*., 2017), either with or without Chiron treatment from 48-72 hours after plating. Gastruloid tissues were dissected for grafting around 120 hours after plating.

### Preparation of Tissues for Grafting

Adherent cells were detached mechanically using a cell scraper in PBS (with calcium and magnesium) to lift intact colonies with minimal sample dissociation. Enzymatic treatments, such as exposure to Accutase^®^, were found to completely preclude cellular NMp-like behaviour after grafting (data not shown). The suspension was transferred to a FBS pre-coated FACS tube and was centrifuged at 170 × *g* for five minutes. The supernatant was discarded and the colonies washed by gentle resuspension in PBS (with calcium and magnesium) before the centrifugation was repeated. The colonies were resuspended in PBS (without calcium and magnesium; Sigma-Aldrich D8537) for labelling with DiI (Thermo Fisher Scientific Vybrant^®^ V22885, 1% v/v.) for 25 minutes in the dark, on ice. The labelled colonies were centrifuged at 170 × *g* for five minutes and the pellet was resuspended in 37°C PBS (with calcium and magnesium) for grafting.

Gastruloid tissues were collected with a micropipette and were dissected into small pieces using a hair loop tool and an eyebrow knife in warm N2B27. Dissected tissues were transferred to an FBS pre-coated FACS tube and were labelled as above.

Explants of embryonic tissue from a square region around the node were dissected with a tungsten needle or an eyebrow knife and were labelled as above.

### Grafting Procedure

Any embryos that were developing abnormally or had flooded with albumen were discarded prior to grafting. A drop of Pannett-Compton saline was pipetted onto the surface of the embryo and two labelled fragments were transferred into the droplet with a mouth pipette. An eyebrow knife tool or an electrolytically sharpened tungsten needle (Brady, 1965) was used to make a small opening in the ectoderm caudal and lateral to the node on each side of the midline. The labelled fragment was positioned in this opening using the tool and the droplet of saline was aspirated to remove any ungrafted labelled cells. The lid of each culture dish was sealed with albumen and the culture was returned to the incubator to heal briefly prior to imaging. Every culture was imaged (see below) within an hour of grafting and approximately 18 hours of grafting; a subset of six embryos was also imaged overnight at 20 minute intervals with time-lapse microscopy in each experiment.

### Microscopy

Widefield, single time points and time-lapse images were acquired with a Zeiss AxioObserver Z1 (Carl Zeiss, UK) using a 5x objective in a humidified 37°C incubator, with the embryo cultures positioned on the inverted lid of a six-well plate. An LED white light illumination system (Laser2000, Kettering, UK) and a Filter Set 45 filter cube (Carl Zeiss, UK) was used to visualise red fluorescence. Emitted light was recorded using a back-illuminated iXon888 Ultra EMCCD (Andor, UK) and the open source Micro-Manager software (Vale Lab, UCSF, USA).

### Quantification

The open-source FIJI ImageJ platform (Schindelin *et al*., 2012) and the pairwise stitching plugin (Preibisch, Saalfeld and Tomancak, 2009) were used for image analysis. Any embryos that were developing abnormally or where the grafted cells had become lost were excluded from further analysis. Each set of images was scored for size and starting position of each graft in relation to the medio-caudal limit of the node, the tissues to which the labelled cells contributed and the final distance between the most rostral and most caudal cells one one side of the midline at the endpoint (around 18 hours after grafting).

Measurements were compiled in Microsoft Excel and were plotted in Python 2.0 with the open source Project Jupyter iPython Notebook and the Matplotlib library. Box plots were prepared in R using the BoxPlotR webtool (Tyers and Rappsilber Labs, Université de Montréal, Canada and University of Edinburgh, UK; respectively), following the conventions described in the accompanying primer (Krzywinski and Altman, 2014). Venn diagrams were produced with the Matplotlib-venn 0.11.5 package.

## Acknowledgements

Thanks are due to Dr David Turner for his assistance in developing the image analysis pipeline and to Professor Alfonso Martinez Arias for his role in devising the study.

## Author Contributions

P.B.-J. and B.S. devised the study and wrote the manuscript. P.B.-J. O.V. and B.S. conducted the experiments. P.B.-J. acquired, quantified and analysed the imaging data.

## Supplementary Figure Captions

**Supplementary Figure 1: Tissue Grafts Extend Many Times their Original Length after Transplantation.** The starting diameter of the grafted tissues was measured in the rostro-caudal direction and is plotted on the *x* axis in (A-C). The final distance between the most rostral and most caudal labelled cells (the axial extent) is shown on the *y* axis in (A-C). (A) Points coloured according to the thresholds (dashed lines) used to score short (<500μm), medium (500–1750μm) and long (>1750μm) rostro-caudal contributions. (B) Points coloured according to whether labelled cells were found in neural tissues only (purple), mesodermal tissues only (green) or both tissue compartments (orange). (C) Points coloured according to whether labelled cells were found in the region associated with the node (red) or not (grey). Grafts that produced medium to long rostro-caudal contributions frequently included both neural and mesodermal tissues and the region around the node.

**Supplementary Figure 2: Fate Mapping Defines the Embryonic NMp-Containing Region Caudal and Lateral to the Node.** Small regions of the embryonic tissues were labelled with Dil; the position of each label is represented as absolute axial and lateral displacements from the caudal limit of the node (at 0,0), as in other figures. The positions of all labels are shown on one plot (top left in (A) and (B)) and separated into different host stages (7.67 represents HH Stage 8^-^). (A) Tissue labels coloured by the rostro-caudal length of their final contributions (blue, <500μm; orange, 500–1750μm; red, >1750μm). (B) Tissue labels coloured by the tissues in which labelled cells are scored after incubation. LPM: Lateral Plate Mesoderm, PSM: Pre-Somitic Mesoderm, N: Node, NT: Neural Tube, SM: Somitic Mesoderm.

**Supplementary Figure 3: Explanted Chick Node Tissue.** Explants of chick node tissue were labelled with Dil and grafted into host embryos; the position of each graft is represented as absolute axial and lateral displacements from the caudal limit of the node (at 0,0), as in other figures. The positions of all grafts are shown on one plot (top left in (A) and (B)) and separated into different host stages (7.67 represents HH Stage 8^-^). (A) Tissue labels coloured by the rostro-caudal length of their final contributions (blue, <500μm; orange, 500–1750μm; red, >1750μm). (B) Tissue labels coloured by the tissues in which labelled cells are scored after incubation. LPM: Lateral Plate Mesoderm, PSM: Pre-Somitic Mesoderm, N: Node, NT: Neural Tube, SM: Somitic Mesoderm.

**Supplementary Figure 4: Self-Renewing Embryonic Stem Cells.** Colonies of ES cells from ES+LIF culture (see Materials & Methods) were labelled with Dil and grafted into host embryos; the position of each graft is represented as absolute axial and lateral displacements from the caudal limit of the node (at 0,0), as in other figures. The positions of all grafts are shown on one plot (top left in (A) and (B)) and separated into different host stages (7.67 represents HH Stage 8^-^). (A) Tissue labels coloured by the rostro-caudal length of their final contributions (blue, <500μm; orange, 500–1750μm; red, >1750μm). (B) Tissue labels coloured by the tissues in which labelled cells are scored after incubation. LPM: Lateral Plate Mesoderm, PSM: Pre-Somitic Mesoderm, N: Node, NT: Neural Tube, SM: Somitic Mesoderm.

**Supplementary Figure 5: Embryonic Stem Cell-Derived NMps, Turner *et al*. Protocol.** Colonies of ES cells differentiated according to the protocol used by Turner *et al*. were labelled with Dil and grafted into host embryos; the position of each graft is represented as absolute axial and lateral displacements from the caudal limit of the node (at 0,0), as in other figures. The positions of all grafts are shown on one plot (top left in (A) and (B)) and separated into different host stages (7.67 represents HH Stage 8^-^). (A) Tissue labels coloured by the rostro-caudal length of their final contributions (blue, <500μm; orange, 500–1750μm; red, >1750μm). (B) Tissue labels coloured by the tissues in which labelled cells are scored after incubation. LPM: Lateral Plate Mesoderm, PSM: Pre-Somitic Mesoderm, N: Node, NT: Neural Tube, SM: Somitic Mesoderm.

**Supplementary Figure 6: Embryonic Stem Cell-Derived NMps, Gouti *et al*. Protocol.** Colonies of ES cells differentiated according to the protocol used by Gouti *et al*. were labelled with Dil and grafted into host embryos; the position of each graft is represented as absolute axial and lateral displacements from the caudal limit of the node (at 0,0), as in other figures. The positions of all grafts are shown on one plot (top left in (A) and (B)) and separated into different host stages (7.67 represents HH Stage 8^-^). (A) Tissue labels coloured by the rostro-caudal length of their final contributions (blue, <500μm; orange, 500–1750μm; red, >1750μm). (B) Tissue labels coloured by the tissues in which labelled cells are scored after incubation. LPM: Lateral Plate Mesoderm, PSM: Pre-Somitic Mesoderm, N: Node, NT: Neural Tube, SM: Somitic Mesoderm.

**Supplementary Figure 7: Chiron-Treated Gastruloid Tissues, Anterior Explants.** Anterior explants of Chiron-treated gastruloid tissues were labelled with DiI and grafted into host embryos; the position of each graft is represented as absolute axial and lateral displacements from the caudal limit of the node (at 0,0), as in other figures. The positions of all grafts are shown on one plot (top left in (A) and (B)) and separated into different host stages (7.67 represents HH Stage 8^-^). (A) Tissue labels coloured by the rostro-caudal length of their final contributions (blue, <500μm; orange, 500–1750μm; red, >1750μm). (B) Tissue labels coloured by the tissues in which labelled cells are scored after incubation. LPM: Lateral Plate Mesoderm, PSM: Pre-Somitic Mesoderm, N: Node, NT: Neural Tube, SM: Somitic Mesoderm.

**Supplementary Figure 8: Chiron-Treated Gastruloid Tissues, Posterior Explants.** Posterior explants of Chiron-treated gastruloid tissues were labelled with DiI and grafted into host embryos; the position of each graft is represented as absolute axial and lateral displacements from the caudal limit of the node (at 0,0), as in other figures. The positions of all grafts are shown on one plot (top left in (A) and (B)) and separated into different host stages (7.67 represents HH Stage 8^-^). (A) Tissue labels coloured by the rostro-caudal length of their final contributions (blue, <500μm; orange, 500–1750μm; red, >1750μm). (B) Tissue labels coloured by the tissues in which labelled cells are scored after incubation. LPM: Lateral Plate Mesoderm, PSM: Pre-Somitic Mesoderm, N: Node, NT: Neural Tube, SM: Somitic Mesoderm.

**Supplementary Figure 9: N2B27-Cultured Gastruloid Tissues.** Explants of gastruloids grown continuously in N2B27 were labelled with DiI and grafted into host embryos; the position of each graft is represented as absolute axial and lateral displacements from the caudal limit of the node (at 0,0), as in other figures. The positions of all grafts are shown on one plot (top left in (A) and (B)) and separated into different host stages (7.67 represents HH Stage 8^-^). (A) Tissue labels coloured by the rostro-caudal length of their final contributions (blue, <500μm; orange, 500–1750μm; red, >1750μm). (B) Tissue labels coloured by the tissues in which labelled cells are scored after incubation. LPM: Lateral Plate Mesoderm, PSM: Pre-Somitic Mesoderm, N: Node, NT: Neural Tube, SM: Somitic Mesoderm.

**Supplementary Figure 10: All Populations Tested Can Produce Long Neural and Mesodermal Contributions, Except Anterior Chiron-Treated Gastruloid Explants.** The distance between the most rostal and most caudal labelled cells from each graft (its axial length) was measured and is plotted as open circles. Box plots show the distribution of axial lengths for each starting population. Centre lines show the medians; crosses denote the sample means. Box limits indicate the 25^th^ and 75^th^ percentiles as determined by the BoxPlotR webtool (see Materials & Methods); whiskers extend 1.5 times the interquartile range (Tukey style). The width of the boxes is proportional to the square root of the sample size.

## References

Baillie-Johnson, P. (2017) The Generation of a Candidate Axial Precursor in Three Dimensional Aggregates of Mouse Embryonic Stem Cells. University of Cambridge. doi: 10.17863/CAM. 13742.

Brady, J. (1965) ‘A simple technique for making very fine, durable dissecting needles by sharpening tungsten wire electrolytically’, Bulletin of the World Health Organization, 32(1), pp. 143–144.

Brown, J. M. and Storey, K. G. (2000) ‘A region of the vertebrate neural plate in which neighbouring cells can adopt neural or epidermal fates’, Current Biology, 10(14), pp. 869–872. doi: 10.1016/S0960-9822(00)00601-1.

Cambray, N. and Wilson, V. (2002) ‘Axial progenitors with extensive potency are localised to the mouse chordoneural hinge.’, Development (Cambridge, England), 129(20), pp. 4855–66. Available at: http://www.ncbi.nlm.nih.gov/pubmed/12361976.

Cambray, N. and Wilson, V. (2007) ‘Two distinct sources for a population of maturing axial progenitors.’, Development (Cambridge, England), 134(15), pp. 2829–40. doi: 10.1242/dev.02877.

Catala, M., Teillet, M. A., de Robertis, E. M. and Le Douarin, N. M. (1996) ‘A spinal cord fate map in the avian embryo: while regressing, Hensen’s node lays down the notochord and floor plate thus joining the spinal cord lateral walls.’, Development (Cambridge, England), 122(9), pp. 2599–610. doi: 10.1016/0896-6273(93)90227-i.

Edri, S., Hayward, P., Baillie-Johnson, P., Steventon, B. and Martinez Arias, A. (2018) ‘An Epiblast Stem Cell derived multipotent progenitor population for axial extension’, *bioRxiv*. doi: 10.1101/242461.

Fehling, H. J., Lacaud, G., Kubo, A., Kennedy, M., Robertson, S., Keller, G. and Kouskoff, V. (2003) ‘Tracking mesoderm induction and its specification to the hemangioblast during embryonic stem cell differentiation.’, Development (Cambridge, England), 130(17), pp. 4217–4227. doi: 10.1242/dev.00589.

Gouti, M., Delile, J., Stamataki, D., Wymeersch, F. J., Huang, Y., Kleinjung, J., Wilson, V. and Briscoe, J. (2017) ‘A gene regulatory network balances neural and mesoderm specification during vertebrate trunk development’, Developmental Cell. Elsevier Inc., In press, pp. 1–19. doi: 10.1016/j.devcel.2017.04.002.

Gouti, M., Tsakiridis, A., Wymeersch, F. J., Huang, Y., Kleinjung, J., Wilson, V. and Briscoe, J. (2014) ‘In Vitro Generation of Neuromesodermal Progenitors Reveals Distinct Roles for Wnt Signalling in the Specification of Spinal Cord and Paraxial Mesoderm Identity.’, PLoS biology, 12(8), p. e1001937. doi: 10.1371/journal. pbio. 1001937.

Hamburger, V. and Hamilton, H. (1993) ‘A Series of Normal Stages in the Development of the Chick Embryo (Reprinted from J. Morphol., 88:1 (1951))’, Developmental Dynamics, 195(1), pp. 231–272.

Henrique, D., Abranches, E., Verrier, L. and Storey, K. G. (2015) ‘Neuromesodermal progenitors and the making of the spinal cord’, Development, 142(17), pp. 2864–2875. doi: 10.1242/dev.119768.

Hooper, M., Hardy, K., Handyside, a, Hunter, S. and Monk, M. (1987) ‘HPRT-deficient (Lesch-Nyhan) mouse embryos derived from germline colonization by cultured cells.’, Nature, 326(6110), pp. 292–295. doi: 10.1038/326292a0.

Krzywinski, M. and Altman, N. (2014) ‘Points of Significance: Visualizing samples with box plots’, Nature Methods. Nature Publishing Group, 11(2), pp. 119–120. doi: 10.1038/nmeth.2813.

McGrew, M. J., Sherman, A., Lillico, S. G., Ellard, F. M., Radcliffe, P. a, Gilhooley, H. J., Mitrophanous, K. a, Cambray, N., Wilson, V. and Sang, H. (2008) ‘Localised axial progenitor cell populations in the avian tail bud are not committed to a posterior Hox identity.’, Development (Cambridge, England), 135(13), pp. 2289–2299. doi: 10.1242/dev.022020.

New, D. A. (1955) ‘A New Technique for the Cultivation of the Chick Embryo in vitro’, Journal of embryology and experimental morphology, 3(December), p. 320.

Papanayotou, C., Benhaddou, A., Camus, A., Perea-Gomez, A., Jouneau, A., Mezger, V., Langa, F., Ott, S., Sabéran-Djoneidi, D. and Collignon, J. (2014) ‘A Novel Nodal Enhancer Dependent on Pluripotency Factors and Smad2/3 Signaling Conditions a Regulatory Switch During Epiblast Maturation’, PLoS Biology, 12(6). doi: 10.1371/journal.pbio. 1001890.

Preibisch, S., Saalfeld, S. and Tomancak, P. (2009) ‘Globally optimal stitching of tiled 3D microscopic image acquisitions’, Bioinformatics, 25(11), pp. 1463–1465. doi: 10.1093/bioinformatics/btp184.

Schindelin, J., Arganda-Carreras, I., Frise, E., Kaynig, V., Longair, M., Pietzsch, T., Preibisch, S., Rueden, C., Saalfeld, S., Schmid, B., Tinevez, J.-Y., White, D. J., Hartenstein, V., Eliceiri, K., Tomancak, P. and Cardona, A. (2012) ‘Fiji: an open-source platform for biological-image analysis’, Nature Methods, 9(7), pp. 676–682. doi: 10.1038/nmeth.2019.

Selleck, M. A., and Stern, C. D. (1991) ‘Fate mapping and cell lineage analysis of Hensen’s node in the chick embryo.’, Development (Cambridge, England), 112(2), pp. 615–26. Available at: http://www.ncbi.nlm.nih.gov/pubmed/1794328.

Steventon, B. and Arias, A. M. (2016) ‘Evo-engineering and the Cellular and Molecular Origins of the Vertebrate Spinal Cord’, Developmental Biology. Elsevier, (January), pp. 0–1. doi: 10.1101/068882.

Tsakiridis, A., Huang, Y., Blin, G., Skylaki, S., Wymeersch, F., Osorno, R. Economou, C., Karagianni, E., Zhao, S., Lowell, S. and Wilson, V. (2014) ‘Distinct Wnt-driven primitive streak-like populations reflect in vivo lineage precursors.’, Development (Cambridge, England), 141(6), pp. 1209–21. doi: 10.1242/dev.101014.

Turner, D. A., Girgin, M., Alonso-Crisostomo, L., Trivedi, V., Baillie-Johnson, P., Glodowski, C. R., Hayward, P. C., Collignon, J., Gustavsen, C., Serup, P., Steventon, B., Lutolf, M. and Martinez, A. A. (2017) Anteroposterior polarity and elongation in the absence of extraembryonic tissues and spatially localised signalling in Gastruloids, mammalian embryonic organoids, Development, doi: 10.1242/dev. 150391.

Turner, D. A., Hayward, P. C., Baillie-Johnson, P., Rue, P., Broome, R., Faunes, F. and Martinez Arias, A. (2014) ‘Wnt/β-catenin and FGF signalling direct the specification and maintenance of a neuromesodermal axial progenitor in ensembles of mouse embryonic stem cells’, Development, 141(22), pp. 4243–4253. doi:10.1242/dev. 112979.

Tzouanacou, E., Wegener, A., Wymeersch, F. J., Wilson, V. and Nicolas, J.-F. (2009) ‘Redefining the progression of lineage segregations during mammalian embryogenesis by clonal analysis.’, Developmental cell, 17(3), pp. 365–76. doi: 10.1016/j.devcel.2009.08.002.

Voiculescu, O., Papanayotou, C. and Stern, C. D. (2008) ‘Spatially and temporally controlled electroporation of early chick embryos’, Nat Protoc, 3(3), pp. 419–426. doi: nprot.2008.10 {pii}\r10.1038/nprot.2008.10.

Wymeersch, F. J., Huang, Y., Blin, G., Cambray, N., Wilkie, R., Wong, F. C. and Wilson, V. (2016) ‘Position-dependent plasticity of distinct progenitor types in the primitive streak.’, eLife, 5(10042), pp. 1–28. doi: 10.7554/eLife.10042.

